# Early maturational emergence of adult-like emotional reactivity and anxiety after brief exposure to an obesogenic diet

**DOI:** 10.1101/2020.03.03.975789

**Authors:** Julio D. Vega-Torres, Matine Azadian, Raul Rios-Orsini, Arsenio L. Reyes-Rivera, Perla Ontiveros-Angel, Johnny D. Figueroa

**Affiliations:** Center for Health Disparities and Molecular Medicine and Department of Basic Sciences, Physiology Division, Department of Basic Sciences, Loma Linda University School of Medicine, Loma Linda, California, USA; San Juan Bautista School of Medicine, Caguas, Puerto Rico; Stanford University School of Medicine, Stanford, California, USA

**Keywords:** PTSD, diet-induced obesity, adolescence, anxiety, fear-potentiated startle, CRHR1

## Abstract

**Background:** Emerging evidence demonstrates that diet-induced obesity disrupts corticolimbic circuits underlying emotional regulation. Studies directed at understanding how obesity alters brain and behavior are easily confounded by a myriad of complications related to obesity. This study investigated the early neurobiological stress response triggered by an obesogenic diet. Furthermore, this study directly determined the combined impact of a short-term obesogenic diet and adolescence on critical behavioral and molecular substrates implicated in emotion regulation and stress.

**Methods:** Adolescent (postnatal day 31) or adult (postnatal day 81) Lewis rats were fed for one week with an experimental Western-like high-saturated fat diet (*WD*, 41% kcal from fat) or a matched control diet (*CD*, 13% kcal from fat). We used the acoustic fear-potentiated startle (FPS) paradigm to determine the effects of the WD on cued fear conditioning and fear extinction. We used c-Fos mapping to determine the functional influence of the diet and stress on corticolimbic circuits.

**Results:** We report that one-week WD consumption was sufficient to induce fear extinction deficits in adolescent rats, but not in adult rats. We identify fear-induced alterations in corticolimbic neuronal activation and demonstrate increased prefrontal cortex CRHR1 mRNA levels in the rats that consumed the WD.

**Conclusions:** Our findings demonstrate that short-term consumption of an obesogenic diet during adolescence heightens behavioral and molecular vulnerabilities associated with risk for anxiety and stress-related disorders. Given that fear extinction promotes resilience, and that fear extinction principles are the foundation of psychological treatments for PTSD, understanding how obesogenic environments interact with the adolescent period to affect the acquisition and expression of fear extinction memories is of tremendous clinical relevance.

**HIGHLIGHTS:**

- Short-term WD consumption during adolescence impairs cued fear extinction memory retention in a fear-potentiated startle paradigm.
- Short-term WD consumption during adolescence attenuates neuronal activation to electric footshock stress in the basomedial nuclei of the amygdala.
- Short-term WD consumption increases CRHR1 mRNA levels in the medial prefrontal cortex.
- Adult LEW rats exhibit increased basal HPA axis tone and heightened emotional reactivity to footshock stress relative to adolescent rats.

## 1. INTRODUCTION

Early-life trauma is linked to obesity and the consumption of obesogenic diets (Duncan et al., 2015; Ehlert, 2013; Farr et al., 2015; Masodkar et al., 2016; Mason et al., 2017; Pagoto et al., 2012; Perkonigg et al., 2009; Roenholt et al., 2012; Wolf et al., 2017). The high co-morbidity between obesity and post-traumatic stress disorders (PTSD) suggest that adaptations to trauma may increase the risk for the consumption of obesogenic diets as a result of the traumatic experience (Godfrey et al., 2018; Kalyan-Masih et al., 2016; Michopoulos et al., 2016). There is also mounting evidence that exposure to obesogenic diets rich in saturated fat foods and sugars have a direct adverse effect on emotional regulation, anxiety-like behaviors, and neural substrates implicated with stress (Baker and Reichelt, 2016; Boitard et al., 2015; Kalyan-Masih et al., 2016; Ortolani et al., 2011; Reichelt et al., 2015; Sivanathan et al., 2015; Vega-Torres et al., 2018). Therefore, early-life exposure to obesogenic diets may predispose individuals to maladaptive stress responses, resulting in increased PTSD risk.

Several lines of evidence suggest that alterations in attention, memory, and learning contribute to the etiology and maintenance of PTSD symptoms (Liberzon and Abelson, 2016; Lissek and van Meurs, 2015). Interestingly, while fear learning emerges early in life, fear memories undergo dynamic changes during adolescence (Baker and Richardson, 2015; Ganella et al., 2017; King et al., 2014). Studies indicate that extinction learning is blunted during adolescence (Pattwell et al., 2012), which has important implications for PTSD treatment. The fear-potentiated startle (FPS) represents a proven and reliable method for examining conditioned fear responses (Davis et al., 1993). This method shows notable face validity, construct validity, and predictive validity in the assessment of behaviors and circuits implicated in PTSD. The highly conserved corticolimbic circuit is critical for cue-elicited fear responses and safety learning (Likhtik and Paz, 2015; Likhtik et al., 2014). In particular, the corticolimbic pathway connecting the medial prefrontal cortex (mPFC) and the amygdala, undergoes dramatic structural reorganization during adolescence (Arruda-Carvalho et al., 2017; Casey, 2015; Silvers et al., 2017) and remains the focus of our recent investigations. Our studies indicate that this corticolimbic pathway is highly vulnerable to the disruptive effects of an obesogenic diet when exposure to the diet starts during adolescence (Vega-Torres et al., 2018). However, without a cohort of animals exposed to the obesogenic diet only during adulthood in our previous studies, it is difficult to determine if adolescence represents a period of vulnerability to the detrimental effects of an obesogenic diet on learned fear and emotional reactivity. Furthermore, few studies have tested the hypothesis that short-term consumption of an obesogenic diet is sufficient to alter fear responses.

This study tested several hypotheses. First, we predicted impaired fear extinction in adolescent rats that consumed an obesogenic diet for one week. Not only would this represent a replication of our previous report in adult rats (Vega-Torres et al., 2018), but it would also extend that finding to diet effects independent of obesity-related processes. Second, we reasoned that short-term exposure to an obesogenic diet would reduce the neuronal activation in corticolimbic regions implicated in fear extinction and anxiolytic effects. Finally, given the robust effects of obesogenic diets on the HPA axis and dopamine systems (Boitard et al., 2015; Khazen et al., 2019; Reyes, 2012; Sharma et al., 2013), and the modulatory actions of these factors on mPFC-amygdala circuit function (Fadok et al., 2009b; Jovanovic et al., 2010; 2020; Veer et al., 2012; Whittle et al., 2016), we hypothesized that the obesogenic diet would increase the HPA tone while reducing dopamine receptor expression in the mPFC.

This study demonstrates that short exposure to an obesogenic diet is sufficient to impair retention of fear extinction training while altering neurobiological substrates implicated with emotional reactivity. These findings reveal a unique interplay between high saturated fat/high-sugar foods and fear extinction during adolescence, which may prove informative for understanding risk factors implicated in stress-related disorders. More importantly, this study suggests that obesity and the consumption of obesogenic diets may represent a mediator of differential anxiety and stress-related disorders psychotherapy treatment outcomes.

## 2. METHODS

### Animals

All the experiments were performed following protocols approved by the Institutional Animal Care and Use Committee (IACUC) at the Loma Linda University School of Medicine. Adolescent (**ADOL**: postnatal day, PND, 24) and adult (**ADUL**: PND 74) male Lewis rats were purchased from Charles River Laboratories (Portage, MI, USA). The rationale for the use of Lewis rats is based on the relevant vulnerabilities of this strain to post-traumatic stress (Kalyan-Masih et al., 2016; Vega-Torres et al., 2018). Immediately upon arrival, the rats were housed in groups (2 per cage) with free access to food and water. Dietary manipulations commenced at PND 31. Animals were kept in customary housing conditions (21 ± 2 C, relative humidity of 45%, and 12-hour light/dark cycle with lights on at 7:00 AM). The body weights were recorded once a week or daily during the week of behavioral testing. Food consumption was quantified at least twice per week. The rats were never food or water restricted.

### Diets

The standard chow diet (**CHOW**, 4-gm% fat, product #: 7001) was obtained from Teklad Diets (Madison, WI, USA), while the matched low-fat purified control diet (**CD**, 5-gm% fat, product #: F7463) and Western-like high-saturated fat diet (**WD**, 20-gm% fat, product #: F7462) were obtained from Bio-Serv (Frenchtown, NJ, USA). There is an increasing awareness of the impact of diet changes on stress reactivity (Sharma et al., 2013; Vega-Torres et al., 2018) and a need for using adequate matched diets in diet-induced obesity research (Pellizzon and Ricci, 2018). Thus, we decided to incorporate a matched low-fat control diet group along with the standard chow diet group. Given that initially both the CHOW and CD groups had identical biometric and behavioral outcomes, we opted to use the more appropriate CD group as control for the WD group. The macronutrient composition and fatty acid profiles are detailed in **Table 1**.

**Table 1.** Detailed composition of the purified diets. Dietary fat percentage was ∼5% in the CD and ∼20% in the WD. The WD contained higher levels of total saturated fatty acids (9-fold difference) and total monounsaturated fatty acids (3-fold difference) when compared to the CD. The levels of total polyunsaturated fatty acids were similar between CD and WD.

|  | **Control Diet (CD)** | **Western Diet (WD)** |
| --- | --- | --- |
|  | **#F7463** | **#F7462** |
| **Macronutrient (main source)** | **% kcal** | **% kcal** |
| Carbohydrates (corn starch) | 68 | 43 |
| Proteins (casein) | 19 | 16 |
| Fats (milk fat) | 13 | 41 |
| *Total kcal* | 3.7 | 4.6 |
| **Sugars** | **g/kg** | **g/kg** |
| Sucrose | 341 | 340 |
| **Fatty Acid Class** | **g/kg** | **g/kg** |
| Saturated | 20.8 | 121 |
| Monosaturated | 12.7 | 492 |
| Polyunsaturated | 11.3 | 11.5 |
| Ratio Omega-3 to Omega-6 | 0.02 | 0.01 |

### Study Design

Behavior testing sessions involved a 20-30 min acclimation period to the testing facility. Following room acclimation, the rats were placed for 5 min inside the acoustic startle reflex (ASR) enclosure and testing chamber and then, returned to their cages. The next day (ADOL: PND 30; ADUL: PND 80), we measured baseline ASR responses, which were used to generate balanced experimental groups. The ASR-based group matching resulted in an even distribution of rats with similar startle responses in all groups. The rats were divided in the following groups: ADOL + CD (*n* = 12), ADOL + WD (*n* = 12), ADUL + CD (*n* = 12), ADUL + WD (*n* = 12). We conducted a sensitivity power analysis (Cohen, 1992) (two-way ANOVA: diet type and age as factors) using *G*Power* (Faul et al., 2007). Analyses revealed that 12 rats per group are sufficient to detect medium effect sizes (*d* = .41) with power (1 - β) set at .80, and α = .05. The fear-potentiated startle (FPS) paradigm was performed to assess the short-term diet effects on cued fear conditioning and fear extinction learning at PND 38-41 (ADOL group) and PND 88-91 (ADUL group). We measured anxiety-like responses in the elevated plus maze (EPM) at PND 42 (ADOL) and PND 92 (ADUL). All the rats were euthanized 48 h following EPM testing. The rats were allowed to consume the custom diets until completion of the study at PND 44 (ADOL) and PND 94 (ADUL). **Figure 1** summarizes the timeline of experimental procedures and behavioral tests.

**Figure 1.**
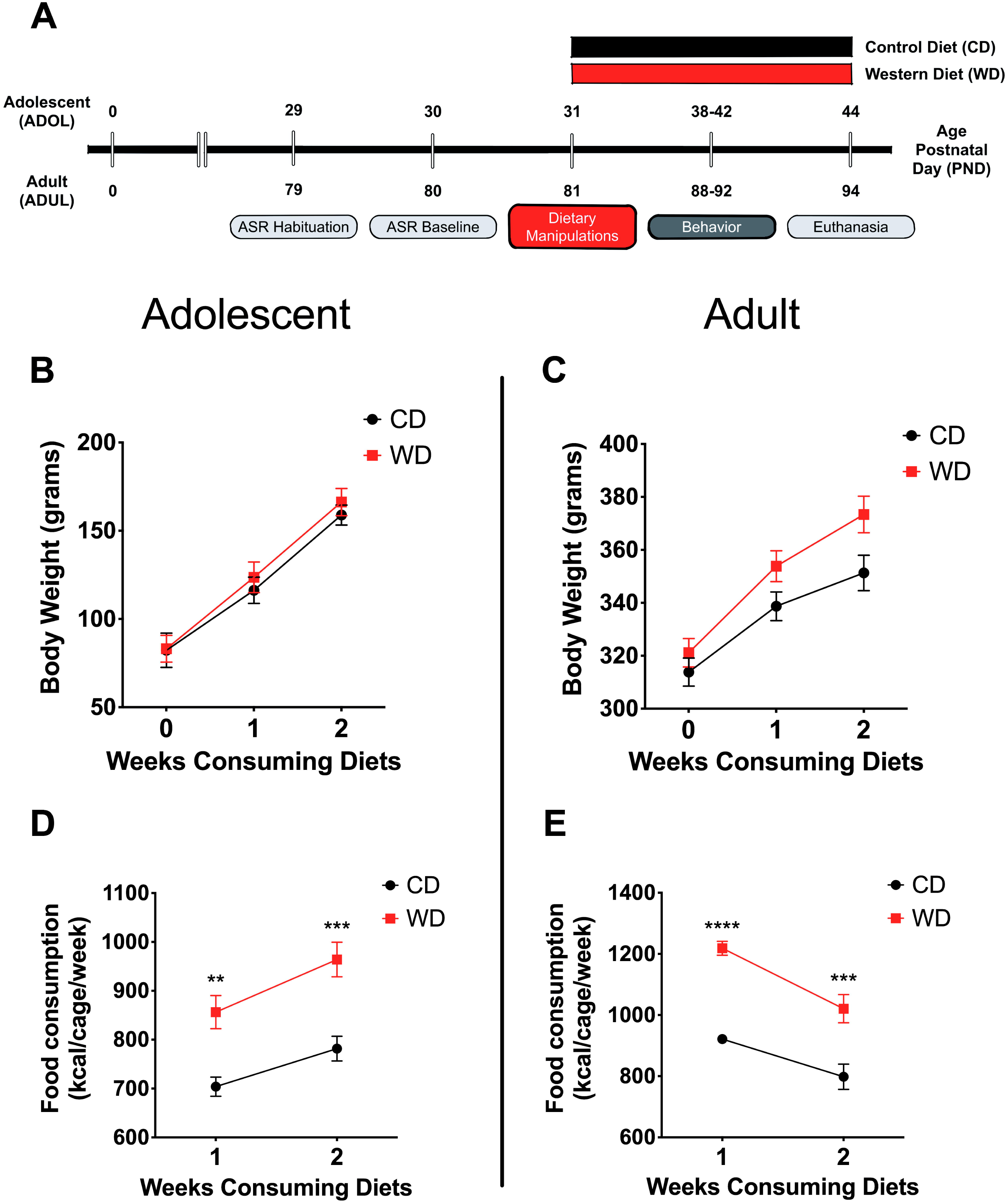
(A) Timeline of experimental procedures. Adolescent (*ADOL*) and adult (*ADUL*) rats were matched based on their acoustic startle reflex (ASR) responses and allocated to one of two diets: control diet (CD) or Western high-saturated fat diet (WD). The rats were exposed to the diets for one week before behavioral testing. A 4-day fear-potentiated startle (FPS) paradigm was used to determine the effects of the WD on cued fear conditioning and fear extinction learning. Additional anxiety-like behaviors were investigated in the elevated plus maze (EPM). All the rats were euthanized one day after the completion of the EPM and brains dissected for RNA extraction. Please refer to the study design in the methods section for more specific technical details on procedures and behavioral tests performed in this study. **Average weekly body weight** in grams for adolescent **(B)** and adult **(C)** groups (diet effect *p* > .05 for both age groups; *n* = 11-12 rats/group). **Average caloric intake** in kilocalories per cage per week for adolescent **(D)** and adult **(E)** groups. WD groups consumed more calories than CD rats, regardless of age (ADOL: diet effect *p* = .002; ADUL: diet effect *p* < .0001; for both groups: *n* = 6 cages/group). Error bars are S.E.M.

### Acoustic startle reflex (ASR)

The ASR experiments were performed using the SR-Lab acoustic chambers (San Diego Instruments, San Diego, CA, USA). ASR magnitudes were measured by placing animals startle enclosures with sensors (piezoelectric transducers) that convert small movements to voltage. Thus, the magnitude of the change in voltage represents the size of the ASR. Acoustic stimuli intensities and response sensitivities were calibrated before commencing the experiments. The ASR protocol has been previously described by our group (Kalyan-Masih et al., 2016; Vega-Torres et al., 2018). Briefly, experimental sessions were 22 min long and started with a 5-min habituation period (background noise = 60 decibels, dB). The rats were then presented with a series of 30 tones (10 tones at each 105 dB) using a 30 sec inter-trial interval (ITI). The acoustic stimuli had a 20 millisecond (ms) duration. Subsequently, the rats were returned to their home cages. Enclosures were thoroughly cleaned and dried following each session. Averaged ASR magnitudes were normalized by weight to eliminate confounding factors associated with body weight (Weight-corrected ASR = maximum startle magnitude in mV divided by body weight at testing day) (Elkin et al., 2006; Gogos et al., 1999; Grimsley et al., 2015; Kalyan-Masih et al., 2016; Vega-Torres et al., 2018). Baseline ASR responses were measured before commencing the dietary manipulations (PND 30: ADOL group and PND 80: ADUL group).

### Fear potentiated startle (FPS)

The fear potentiated startle (FPS) protocol was adapted from Dr. Michael Davis (Davis, 2001) and detailed in our previous studies (Vega-Torres et al., 2018). Each FPS session started with a 5-min acclimation period (background noise = 60 dB). During the first session of the paradigm (fear training), the rats were trained to associate a light stimulus (conditioned stimulus, CS) with a 0.6 mA footshock (unconditioned stimulus, US). The conditioning session involved 10 CS + US pairing presentations. During each CS + US presentation, the light (3200 ms duration) was paired with a co-terminating footshock (500 ms duration). Light-shock pairings were presented in a quasi-random manner (ITI = 3-5 min). Cued fear acquisition was measured 24 h later. During the second session (fear learning testing; pre-extinction FPS), the rats were first presented with 15 startle-inducing tones (*leaders*; 5 each at 90 dB, 95 dB, and 105 dB) delivered alone at 30 sec ITI. Subsequently, the rats were presented with 60 test trials. For half of these test trials, a 20 ms tone was presented alone (**tone alone trials**; 10 trials for each tone: 90 dB, 95 dB, and 105 dB). For the other half, the tone was paired with a 3200 ms light (**light + tone trials**; 10 trials for each pairing: 90 dB, 95 dB, and 105 dB). The 60 trials were divided into 10 blocks of 6 test trials each which included 3 **tone alone trials** and **3 light + tone trials**. To conclude the testing session, the rats were presented with 15 startle-inducing tones (*trailers*; 5 each at 90 dB, 95 dB, and 105 dB) delivered at 30 sec ITI. Trials in this session were presented in a quasi-random order (ITI = 30 sec). The startle-eliciting tones had a 20 ms duration. One day after fear conditioning testing, the rats were exposed to a single extinction-training session. The extinction training session consisted of 30 CS alone presentations (light without shock or noise bursts) with a duration of 3700 ms (ITI = 30 sec). One day after fear extinction training, we determined fear extinction acquisition using the same FPS session that was used to measure fear acquisition. It is noteworthy that in this study we shortened the fear extinction training protocol to a single session as opposed to 3 sessions in our previous report in adult rats (Vega-Torres et al., 2018). A single extinction training session enables accurate and precise fear extinction measurements without flooring effects in adolescent Lewis rats. We assessed fear learning and fear extinction learning by comparing the startle amplitude from light + tone trials (conditioned + unconditioned stimulus, CS + US) relative to tone alone trials (unconditioned stimulus, US). FPS data were reported as the proportional change between US and CS + US [%FPS = ((Light + Tone Startle) − (Tone Alone Startle)) / (Tone Alone Startle) × 100] (Walker and Davis, 2002). Fear recovery, a proxy for fear extinction memory retention, was scored as the ratio between the FPS value from fear extinction and fear learning testing sessions [%Fear recovery = (FPS extinction / FPS learning) × 100].

### Elevated Plus Maze (EPM)

The near infrared-backlit (NIR) elevated plus maze (EPM) consisted of two opposite open arms (50.8 × 10.2 x 0.5 cm) and two opposite enclosed arms (50.8 × 10.2 x 40.6 cm) (Med Associates Inc., Fairfax, VT). The arms were connected by a central 10 × 10 cm square-shaped area. The maze was elevated 72.4 cm from the floor. Behaviors were recorded in a completely dark room. The rats were placed on the central platform facing an open arm and allowed to explore the EPM for 5 min. The apparatus was cleaned after each trial (70% ethanol, rinsed with water, and then dried with paper towels). The behaviors were monitored via a monochrome video camera equipped with a NIR filter and recorded and tracked using Ethovision XT (Noldus Information Technology, Leesburg, VA). In rats, changes in the percentage of time spent on the open arms (OA) indicate changes in anxiety (Pellow et al., 1985) and the number of closed arm (CA) entries is the best measure of locomotor activity (File, 1992). These data are used to calculate the anxiety index (Cohen et al., 2012; Contreras et al., 2014; Kalyan-Masih et al., 2016):

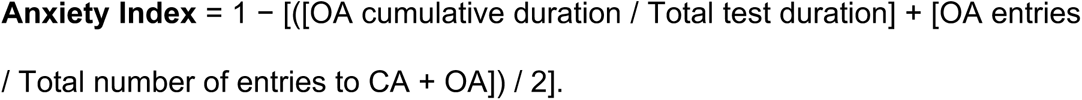

### c-Fos Free-Floating Immunofluorescence

A separate cohort of adolescent and adult rats was exposed to electric footshock stress using the same paradigm employed during fear conditioning. This session consisted of 10 CS + US pairing presentations with a light (3200 ms duration) paired with a co-terminating footshock (0.6 mA footshock; 500 ms duration). Light-shock pairings were presented in a quasi-random manner (ITI = 30 s – 2 min). One h after the session, the rats were sacrificed via transcardiac perfusion with 4% paraformaldehyde (PFA) using the Perfusion Two™ Automated Pressure Perfusion System (Leica Biosystems, Buffalo Grove, IL) (*n* = 12; 3 rats per group).

The brains were removed from the cranial vault 4 h after fixation and postfixed in 4% PFA for 24 h. The brains were then washed with phosphate buffered saline (PBS) and cryoprotected with sucrose (30%) for 12-16 h at 4°C prior to embedding in Tissue-Tek® O.C.T.™ compound (Sakura, Torrance, CA). **Tissue Sampling**: Brain tissue was cut coronally at 25 µm thickness. All immunohistochemical techniques were performed on free-floating sections. Based on the Paxinos and Watson rat brain atlas (from Bregma: +3.20 mm to +2.20 mm), 10 sections were chosen at a 75-μm interval (one of every four sections) covering a total 1000 μm of area containing medial prefrontal cortex (Paxinos and Watson, 2006). An additional 13 sections were cut and chosen in the same manner (from Bregma: −2.30 mm to −3.60 mm) covering a total 1300 μm of area containing the amygdala. **c-Fos Immunofluorescence**: The free-floating tissue sections were washed with PBS and then incubated for blocking with permeable buffer (0.3% Triton-X 100 in PBS) containing 10% normal goat serum for 50 min. The sections were incubated overnight at 4°C in the primary antibodies against c-Fos (1:6400; catalog #2250S; Cell Signaling Technology, Danvers, MA), and then incubated with the secondary Alexa Fluor® 448 (1:800; catalog #4412; Cell Signaling Technology). The slices were then rinsed with PBS, mounted on microscope slides with ProLong™ Gold Antifade mountant containing DAPI (Molecular Probes, Eugene, OR) and cover-slipped. **Analyses:** The medial prefrontal cortex (mPFC) and amygdala were identified as region of interest (ROI) based on common histological landmarks (Paxinos and Watson, 2006). We used randomization of location and orientation within navigation windows to sample within the mPFC and amygdala. We sampled from infralimbic and prelimbic cortices of the mPFC. Within the amygdala, the anterior basolateral amygdaloid nucleus (BL), the posterior basomedial amygdaloid nucleus (BM), and the ventromedial lateral amygdaloid nucleus (L) were analyzed. c-Fos-positive cells were counted within an unbiased counting frame using cell count software (Keyence Corp. of America, Itasca, IL). A Keyence Biorevo BZ-9000 All-In-One Fluorescence Microscope (Keyence) equipped with a Nikon CFI Plan Apo λ20X objective (Nikon, Melville, NY) was used to collect *z*-stacks (numerical aperture: .75; working distance: 1.0 mm). Digitalization and image quantification were carried out by blinded observers. Cells identified as c-Fos positive after threshold correction were quantified automatically using the Macro Cell Count Software (batch image analysis tool from Keyence). For immunofluorescence analyses, a minimum of 5 images per area per animal were used (depending on the size of the ROI). Image analyses were averaged per ROI and the total sample number used for statistical analysis equaled the number of animals used.

### Real-time quantitative polymerase chain reaction (real-time qPCR)

Rats were euthanized with Euthasol (Virbac, Fort Worth, TX, USA) and perfused transcardially with phosphate buffer saline (PBS). Following the perfusion procedure, the prefrontal cortex was isolated. Total RNA was extracted using Trizol (Invitrogen Life Technologies, Carlsbad, CA). The only change to the recommended protocol was using 1-bromo-3-chloropropane (BCP, 0.1 mL per 1 mL of Trizol) instead of chloroform. BCP was obtained from Molecular Research Center (Cincinnati, OH). RNA concentration was determined on a NanoDrop spectrophotometer (Thermo Scientific, Waltham, MA). We used 1 microgram of the total RNA for cDNA synthesis (iScript cDNA Synthesis Kit, Cat. #170-8891, Bio-Rad Laboratories, Hercules, California, USA). cDNA synthesis protocol was performed according to manufacturer’s instructions. The total volume of the cDNA synthesis reaction mixture was 20 microliters (4 microliters, iScript reaction mix; 1 microliter, iScript reverse transcriptase; 15 microliters, nuclease-free water and 1 microgram of RNA). After completion of cDNA synthesis, 80 microliters of nuclease-free water were added to dilute the 20 microliters of synthesized cDNA. The cDNA was amplified by PCR using the primer sets described in **Table 2**. Real-time PCR amplification and analyses were carried out on the CFX96 Real-Time PCR Detection System (Bio-Rad Laboratories, Hercules, California, USA). Real-time qPCR conditions were optimized and 25 microliters reactions were prepared. The PCR reactions contained: 12.5 microliters of iQ SYBR Green Supermix (Cat. #170-8882, Bio-Rad Laboratories, Hercules, California, USA), 1 microliter of a mixture of 10 micromolar forward/reverse primer, 6.5 microliters of water, and 5 microliters of the previously synthesized cDNA. The PCR protocol started with 5 minutes at 95°C. This was followed by 40 cycles of: 15 seconds at 95°C for denaturation and 1 minute at 60°C for annealing/extension. The relative levels of mRNA were calculated using the comparative C*t* (crossing threshold). Each sample was normalized to its GAPDH mRNA content. Relative gene expression levels were normalized to the adolescent CD group and expressed as fold change.

**Table 2.**
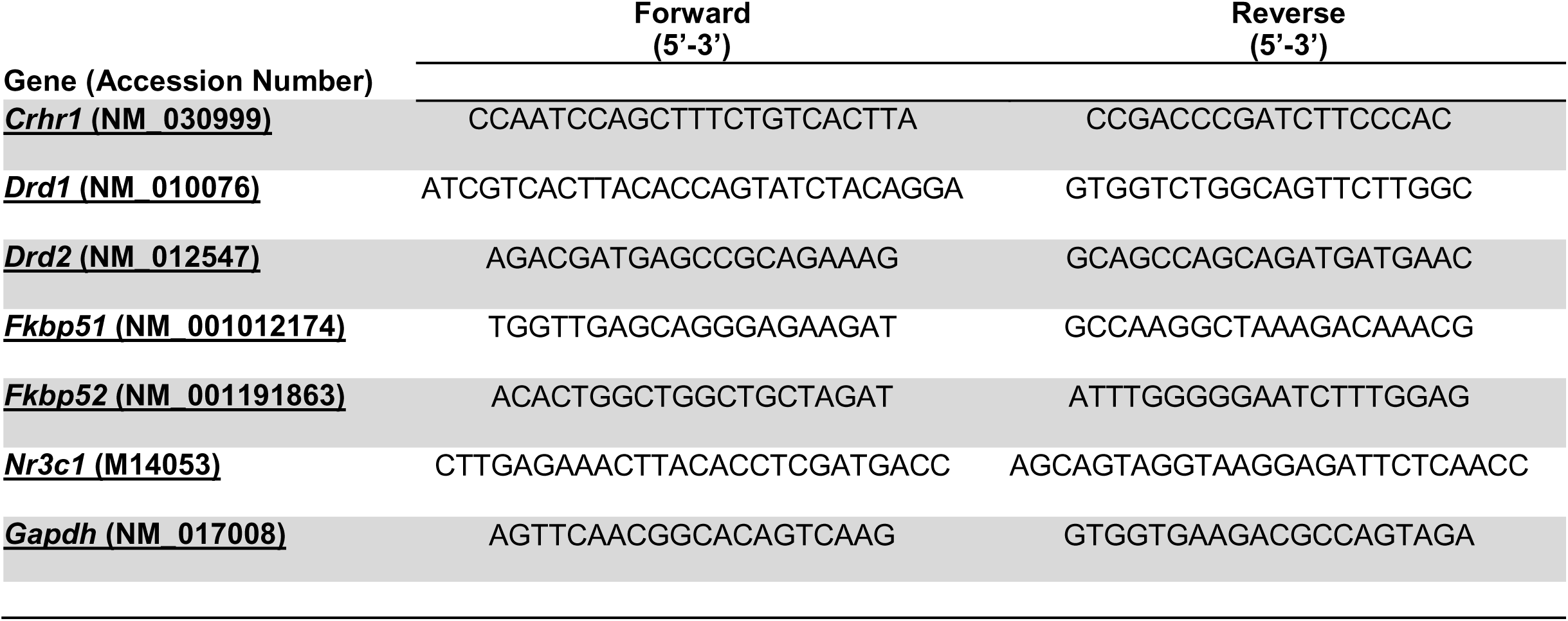
Sequence of primers used for RT-qPCR.

### Glucose Measurements

Blood glucose levels were measured in anesthetized rats by cutting the tail tip before inducing euthanasia. Glucose levels were measured using a glucometer (OneTouch UltraMini, LifeScan).

### Statistical Analysis

We analyzed the data using Graphpad Prism version 8.0. Shapiro-Wilk statistical analyses were used to determine sample distribution. The Brown-Forsythe test was used to test for the equality of group variances. When appropriate, two-way analysis of variance (ANOVA) was used to examine the effect of the diet type, age, and interaction between factors on outcomes measures. Multiple comparisons were made using Dunnett’s (following Welch’s ANOVA) or Sidak’s (repeated measures two-way ANOVA) tests. The ROUT method was used to investigate outliers. We considered differences significant if *p* < .05. The data is shown as the mean ± standard error of the mean (S.E.M.). We also conducted a post-hoc power analysis with the *G*Power* program (Faul et al., 2007). For two-way ANOVA analyses, the statistical power (1-β) for data including the four study groups was .79 for detecting a medium size effect (*d* = .41), whereas the power exceeded .99 for the detection of a large effect size (*d* = .8). This indicates that this study is adequately powered at or above a moderate size level (*d* = .4). Therefore, if chosen at random, the probability that adolescent rats that consumed the WD described here will exhibit alterations in fear-related behaviors relative to controls is .61.

## 4. RESULTS

### Bodyweight and food consumption

This study investigated early brain and behavior responses triggered by a short-term exposure to an obesogenic diet (**Figure 1A**). Body weight and caloric intake were measured weekly. We found that short-term exposure to the obesogenic WD did not alter body weights in adolescent or adult rats. Repeated measures two-way ANOVA revealed a significant effect of week [*F*_(1.21, 25.35)_ = 395.9, *p* < .0001], but no diet [*F*_(1, 21)_ = .24, *p* = .63] or interaction [*F*_(2, 42)_ = .88, *p* = .42] effects on body weight for adolescent rats (**Figure 1B**). Analyses showed a significant effect of week [*F*_(1.42, 31.14)_ = 677.8, *p* < .0001], but no diet [*F*_(1, 22)_ = 3.2, *p* = .09] effects on body weight in adult rats. Interestingly, we found significant interaction between diet and week in adult rats [*F*_(2, 44)_ = 17.91, *p* < .0001] (**Figure 1C**). No significant interaction [*F*_(1, 36)_ = 1.15, *p* = .29] or main effects of the diet [*F*_(1, 36)_ = .04, *p* = .83] and age [*F*_(1, 36)_ = 1.76, *p* = .19] were observed for circulating blood glucose levels at endpoint.

In agreement with previous findings, we found significant changes in caloric intake in the rats that consumed the obesogenic diet. In adolescent rats, we found a significant effect of week [*F*_(1, 10)_ = 58.66, *p* < .0001] and diet [*F*_(1, 10)_ = 17.92, *p* = .002], while no significant interactions [*F*_(1, 10)_ = 1.52, *p* = .25] on caloric intake (**Figure 1D**). Sidak’s post-hoc analyses showed that differences in caloric intake were statistically significant at week 1 (22% increase; *p* = .003). and 2 (23% increase; *p* = .0005) when comparing CD and WD adolescent groups. Similarly, in adult rats, we found a significant effect of week [*F*_(1, 10)_ = 21.29, *p* = .001] and diet [*F*_(1, 10)_ = 62.28, *p* < .0001], while no significant interactions [*F*_(1, 10)_ = 1.15, *p* = .31] on caloric intake (**Figure 1E**). Post-hoc analyses showed that differences in caloric intake were statistically significant at week 1 (32% increase; *p* < .0001) and 2 (28% increase; *p* = .0003) when comparing CD and WD adult groups. The caloric intake of adolescent rats exposed to the WD was similar to that of adult rats (∼900 kcal/cage/week). Notably, while adolescent rats increased caloric intake during week 2 (relative to week 1), the adult rats reduced their caloric intake. This suggests opposite effect of footshock stress on food consumption in adolescent and adult rats undergoing a cued fear paradigm.

### Acute WD exposure attenuates fear extinction learning in adolescent, but not in adult rats

We recently reported that chronic consumption of an obesogenic WD during adolescence leads to impairments in fear-related associative learning and extinction in adult rats (Vega-Torres et al., 2018). However, it remains unclear from our studies whether adolescence and the obesogenic diet are predisposing and precipitating factors for the reported fear impairments. This follow-up study was designed to investigate the effects of short-term exposure to a WD on cued fear conditioning and fear extinction learning before the onset of an obesogenic phenotype in adolescent and adult rats. We used the acoustic startle reflex (ASR) baseline values to assign and match the rats in each group based on their unconditioned responses to acoustic stimulation. Welch’s ANOVA demonstrated no significant differences in baseline weight-corrected ASR responses between groups [*F*_(3, 24.4)_ = .21, *p* = .88] (Figure Not Shown). Following the ASR-based matching, the rats were exposed to either the control or the obesogenic custom diet and fed *ad libitum* for one week before behavioral testing. Anxiety and fear-related responses were investigated using the fear-potentiated startle (FPS) paradigm (**Figure 2A**). The rats were conditioned to learn an aversive association between a light (conditioned stimulus, *CS*) and a foot shock (unconditioned stimulus, *US*) during the first day of the FPS paradigm. ANOVA revealed significantly reduced startle responses to the footshocks in adult rats when compared to adolescent rats [*F*_(3, 24.24)_ = 21.13, *p* < .0001; CD rats: *p* = .0001; WD rats: *p* < .0001] (**Figure 2B**). Acquired fear was tested 24 h after conditioning and defined as significant differences in startle amplitudes between the US (tone alone) and the CS + US (light + tone). All groups showed significant differences between amplitudes of US and CS + US (*p* < .05; data not shown). We calculated the potentiation of the startle as a proxy for fear conditioning and found that one-week WD consumption or age did not alter fear learning [*F*_(3, 19.7)_ = 1.52, *p* = .24] (**Figure 2C**). Fear extinction training and fear extinction testing were performed on days 3 and 4 of the FPS paradigm, respectively. Successful fear extinction is defined as similar startle responses when comparing the US trials to CS + US trials (Vega-Torres et al., 2018). Although FPS responses following extinction testing were similar between groups [*F*_(3, 19.7)_ = 1.78, *p* = .18] (Figure Not Shown), we found heightened fear recovery in the adolescent rats that consumed the WD relative to CD ADOL rats [Welch’s *F*_(3, 18)_ = 6.27, *p* = .004; CD: 14.9 ± 17.92 vs. WD: 123 ± 45.21; *t*_(14.37)_ = 2.22, *p* = .04] (**Figure 2D**). This finding indicates that a short-term exposure to a WD is sufficient to impair fear extinction acquisition and/or retrieval in adolescent rats.

**Figure 2.**
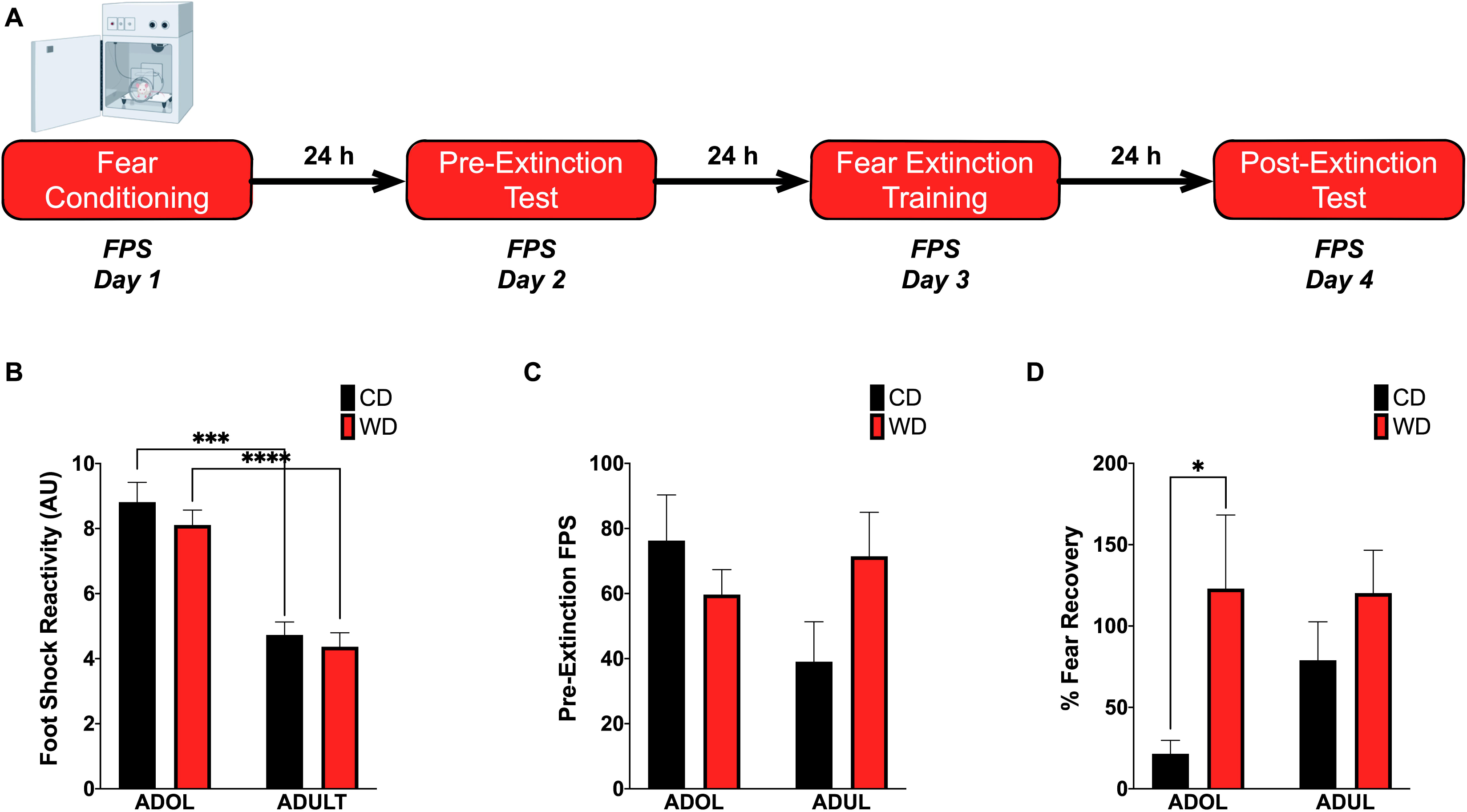
Short-term WD exposure attenuates fear extinction in adolescent rats. **(A)** Fear-potentiated startle (FPS) paradigm. Following habituation and acclimation periods, the rats received a series of light–shock pairings (*FPS day 1, fear conditioning*). Twenty-four h later (*FPS day 2, pre-extinction testing*), the rats were returned to the startle chambers and enclosures and presented with startle stimuli (leaders). Subsequently, rats received startle stimuli presented alone (noise-alone trial) and startle stimuli after onset of the conditioning light (light–noise trials). The rats were presented with startle-eliciting stimuli again at the end of the session (trailers). The two trial types were presented in a balanced mixed order. Twenty-four h later (*FPS day 3, fear extinction training*), the rats were returned to the startle chambers and enclosures and presented with trials of light without shock or noise bursts. One day after fear extinction training (*FPS day 4, post-extinction FPS*), we determined fear extinction acquisition and retention using the same FPS session that was used to measure fear learning. **(B)** Average pre-extinction FPS responses are similar between groups (*p* > .05; *n* = 9-12 rats/group). **(C)** Average post-extinction FPS responses (*p* > .05; *n* = 9-12 rats/group). **(D)** Average percent change in FPS responses between testing sessions as an index of fear recovery following fear extinction training. Adolescent (*ADOL*) WD rats exhibited significantly more FPS than ADOL CD rats, indicating disrupted fear extinction retention (*, *p* < .05; *n* = 9-12 rats/group). Error bars are S.E.M.

### Acute WD exposure enhances startle sensitization in adolescent rats

We reported that chronic consumption of an obesogenic WD during adolescence enhances both pre- and post-extinction background anxiety (BA), as evaluated by changes in ASR responses following cued fear conditioning in adult rats (Vega-Torres et al., 2018). Here, we demonstrate that adult rats exhibited greater background anxiety before extinction training [Welch’s *F*_(3, 23.3)_ = 9.84, *p* = .0002; Dunnett’s post hoc showed an age effect for both the CD (*p = .02*) and WD (*p = .04*) rats] (**Figure 3A**). Notably, we found higher background anxiety in the adolescent rats that consumed the WD relative to CD adolescent rats following fear extinction training [*F*_(3, 21.2)_ = 7.16, *p* = .002; CD ADOL vs. WD ADOL post hoc *p =* .02; CD ADOL vs. CD ADUL post hoc *p =* .04] (**Figure 3B**), supporting an anxiogenic effect of the WD. Short-term habituation of the ASR measures non-associative learning and is determined using the ratio percentage from leader trials to trailer trails within each FPS testing session. We found reduced habituation of the ASR in the adult rats when compared to adolescents before fear extinction training [*F*_(3, 19.7)_ = 18.87, *p* < .0001; Dunnett’s test revealed an age effect for both diet groups CD (*p =* .01) and WD (*p <* .0001)] (**Figure 3C**), suggesting heightened sensitization to footshocks in adult rats. The effect of age in the short-term habituation of the ASR was absent following extinction training [*F*_(3, 22.4)_ = 1.74, *p* = .18] (**Figure 3D**). The long-term habituation of the ASR was determined using the ratio percentage from trailer trials to trailer trails between FPS testing sessions. Analyses revealed that adult rats exhibited reduced long-term habituation of the ASR [*F*_(3, 20.8)_ = 9.58, *p* = .0004; Dunnett’s test revealed an age effect for both diet groups CD (*p =* .02) and WD (*p <* .037)] (Figure Not Shown).

**Figure 3.**
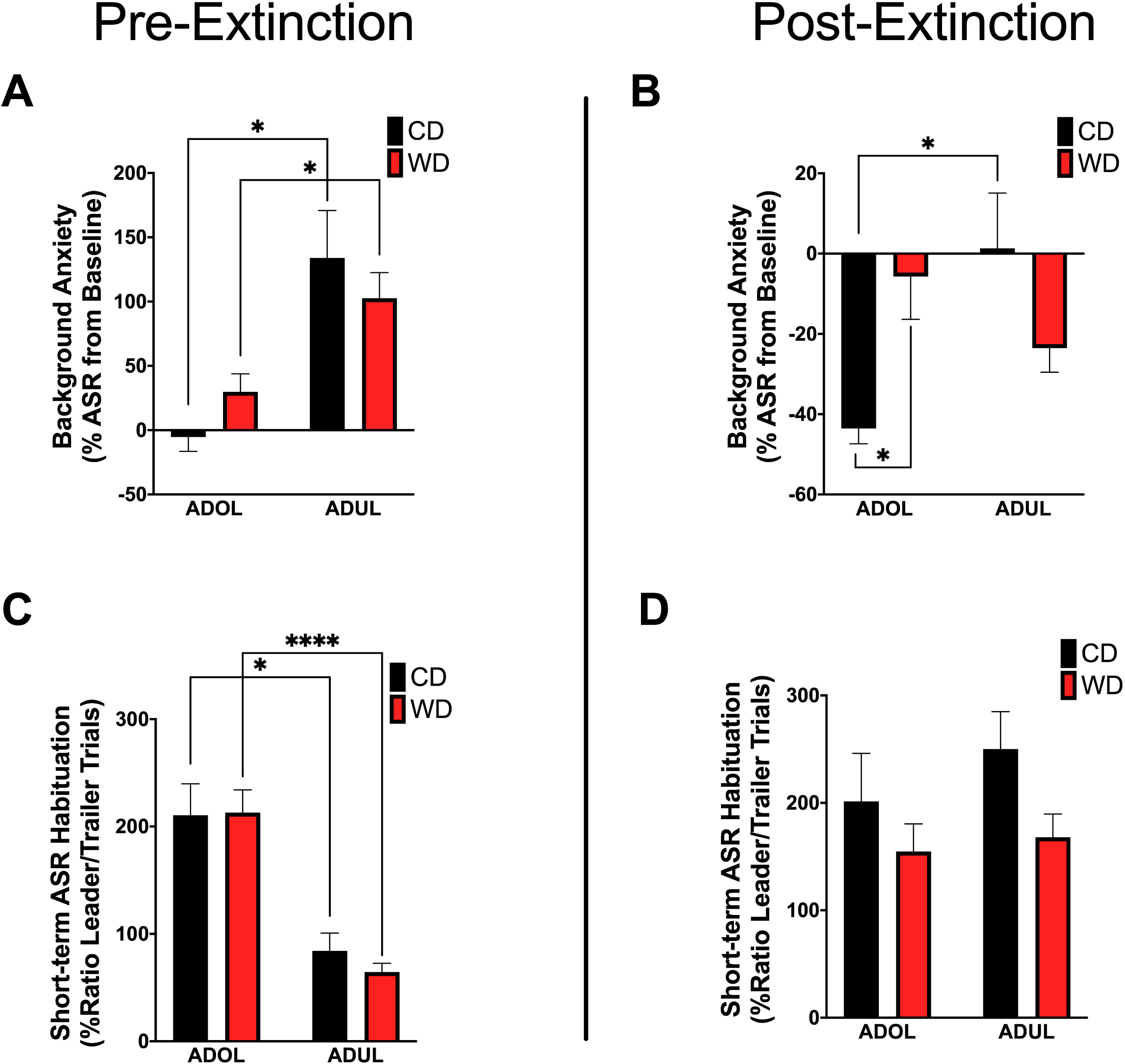
Age and diet influence ASR reactivity and plasticity. **(A)** Average percent change in acoustic startle reflex (ASR) responses following fear conditioning relative to baseline. ADUL rats exhibited higher ASR responses to cued fear conditioning (*, *p* < .05; *n* = 12 rats/group). **(B)** Average percent change in ASR reactivity following fear extinction training. The ADOL rats that consumed the WD exhibited increased indices of anxiety relative to ADOL CD rats (*, *p* < .05; *n* = 12 rats/group). Post-extinction background anxiety in ADOL WD rats was similar to that observed in adult rats (*p* > .05). **(C)** Average pre-extinction short-term habituation of the ASR. ADUL rats exhibited higher within-session ASR responses relative ADOL rats (*, *p* < .05; **, *p* < .01; *n* = 11-12 rats/group). **(D)** All groups exhibited similar short-term habituation of the ASR following fear extinction (*p* > .05; *n* = 10-12 rats/group). Error bars are S.E.M.

### Age increases locomotor activity in the elevated plus maze

Given that footshock stress and obesogenic diet consumption modulate anxiety-like behavior, we carried out analyses of behavioral responses in the elevated plus maze (EPM) one day following fear extinction testing (**Figure 4A**). There was no difference in the duration in the open arms [*F*_(3, 21.6)_ = 1.6, *p* = .21] (**Figure 4B**). The locomotor activity, reflected by the number of crossings between closed arms, was different among groups [*F*_(3, 23.19)_ = 6.51, *p* = .002] (**Figure 4C**). Ambulation was higher in WD ADUL rats than in WD ADOL rats (adjusted *p* = .001). It is noteworthy to mention that we found very robust avoidance responses and anxiety-like behaviors relative to previous studies, suggesting that the fear conditioning paradigm used in this study is anxiogenic. The analysis of the anxiety index, which incorporates several behavioral measures in the maze, revealed no significant differences between groups [*F*_(3, 22.98)_ = 1.72, *p* > .05] (**Figure 4D**), possibly reflecting a ceiling effect.

**Figure 4.**
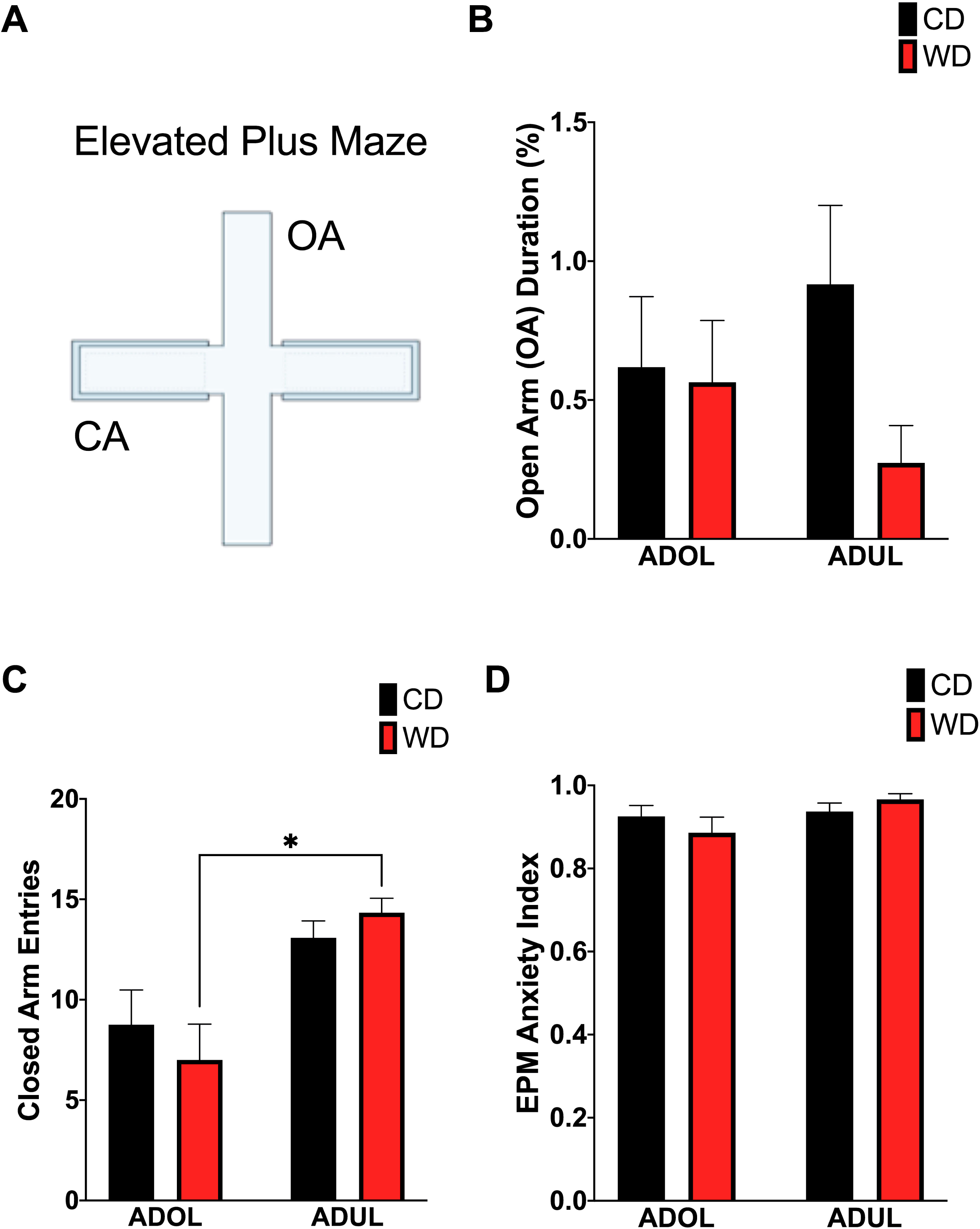
Age influences locomotor activity in the elevated plus maze (EPM). **(A)** Schematic representation of the EPM. **(B)** Average time spent in open arms (expressed as percentage of total time in maze) was similar between groups (*p* > .05; *n* = 10-12 rats/group). **(C)** Average entries to closed arms. ADUL WD rats exhibited higher ambulation when compared to ADOL WD rats (*, *p* < .05; *n* = 10-12 rats/group). **(D)** There was no significant difference in the anxiety index (*p* > .05; *n* = 10-12 rats/group). Error bars are S.E.M.

### Acute WD exposure during adolescence attenuates footshock stress-induced c-Fos expression in the basomedial amygdala

We reported that the consumption of an obesogenic diet has detrimental consequences for the structural integrity of mPFC-amygdala circuits implicated in emotion regulation and fear (Vega-Torres et al., 2018). To further characterize neural substrates impacted by obesogenic diets, we evaluated neuronal activation in mPFC and amygdala regions implicated in fear. We mapped c-Fos expression one hour after the rats were exposed to a cued fear conditioning session. We found no significant main effects of diet [*F*_(1, 8)_ = .61, *p* = .45], age [*F*_(1, 8)_ = .33, *p* = .58], or interactions [*F*_(1, 8)_ = 1.74, *p* = .22] on footshock-induced c-Fos expression in the prelimbic region of the mPFC (**Figure 5A**). Similarly, the diet type [*F*_(1, 8)_ = .06, *p* = .81], age [*F*_(1, 8)_ = 2.06, *p* = .18], and interactions between factors [*F*_(1, 8)_ = 4.55, *p* = .06] did not have a significant effect on c-Fos protein levels in the infralimbic region of the mPFC (**Figure 5B**). The basolateral, basomedial, and lateral amygdaloid nuclei were also assessed for cued fear conditioning-induced c-Fos expression. While analyses revealed no significant effects of the diet, age, and interactions in c-Fos expression in the basolateral (diet: *F*_(1, 8)_ = 1.20, *p* = .31; age: *F*_(1, 8)_ = .11, *p* = .76; interaction: *F*_(1, 8)_ = .23, *p* = .65) (Figure Not Shown) and lateral amygdaloid nuclei (diet: *F*_(1, 8)_ = 1.97, *p* = .20; age: *F*_(1, 8)_ = 3.78, *p* = .09; interaction: *F*_(1, 8)_ = 2.40, *p* = .16) (**Figure 5C**), we found that the obesogenic diet reduced c-Fos expression levels in the basomedial nucleus of the amygdala (diet: *F*_(1, 8)_ = 10.15, *p* = .01; age: *F*_(1, 8)_ = .4, *p* = .55; interaction: *F*_(1, 8)_ = 3.21, *p* = .11] (**Figure 5D**). Sidak’s post hoc analysis revealed that acute exposure to the WD during adolescence led to a significant reduction in basomedial amygdala c-Fos expression relative to control rats (adjusted *p* = .02).

**Figure 5.**
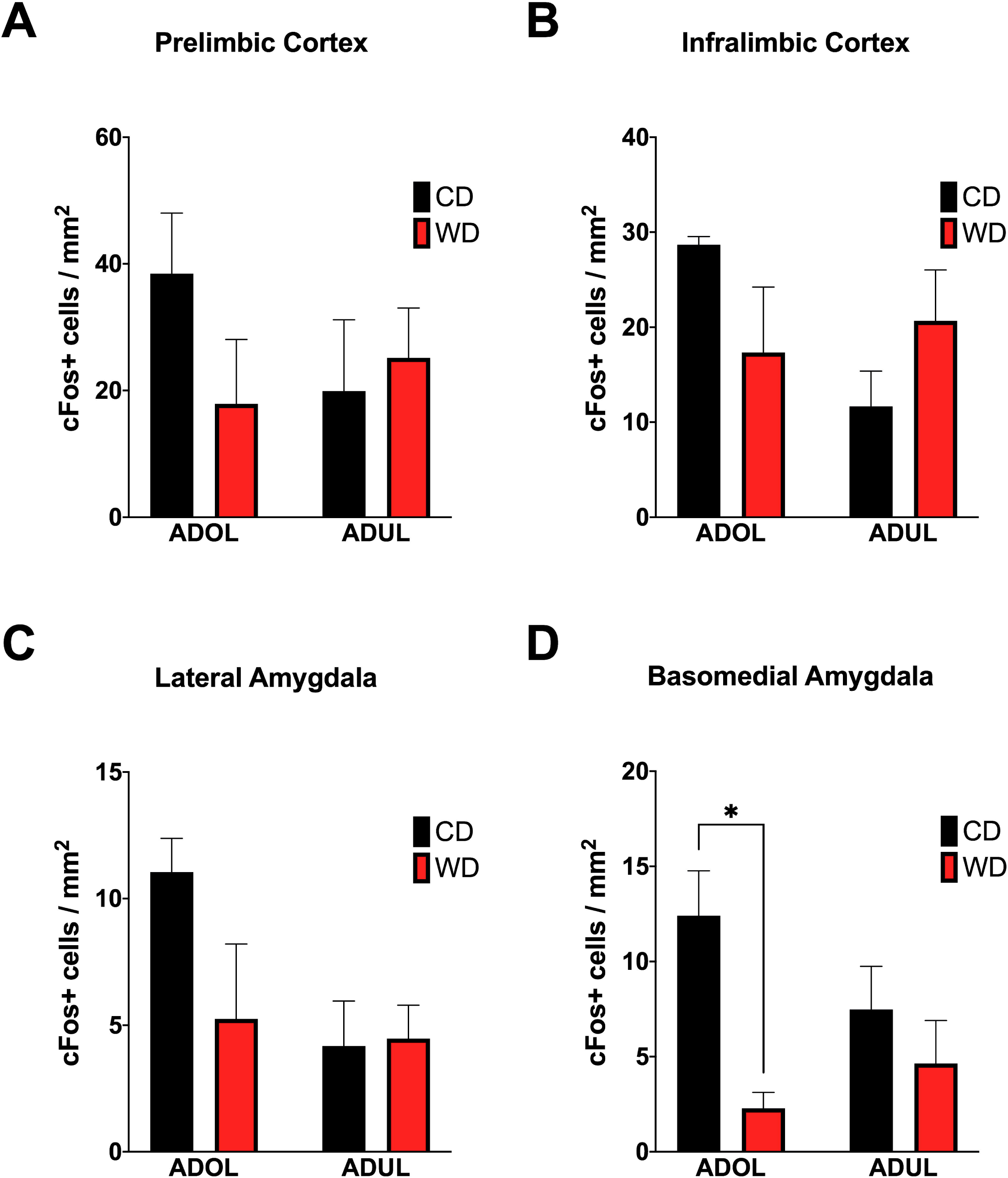
WD consumption during adolescence attenuates neuronal activation to footshock stress in the basomedial nuclei of the amygdala. Average expression of the immediate-early gene c-Fos in prelimbic **(A)** and infralimbic **(B)** medial prefrontal cortex, and in the lateral **(C)** and basomedial **(D)** nuclei of the amygdala. ADOL WD rats showed attenuated footshock-induced c-Fos expression in BMA neurons relative to ADOL CD rats (*, *p* < .05; *n* = 3 rats/group). Error bars are S.E.M.

### Age, short-term obesogenic diet exposure, and interactions between these factors modulate the levels of critical fear-modulating biomarkers

We evaluated the levels of critical fear-associated biomarkers to gain insights on putative early (mal)adaptive mechanisms impacted by a short-term exposure to the obesogenic diet. We found a significant effect of age on plasma corticosterone (CORT) levels (diet: *F*_(1, 24)_ = .007, *p* = .93; age: *F*_(1, 24)_ = 13.22, *p* = .001; interaction: *F*_(1, 24)_ = .51, *p* = .48) (**Figure 6A**). The adult rats exposed to the short-term WD had significantly higher CORT levels when compared to the adolescent rats that consumed the same diet (*p = .01*). Consistent with this finding, age had a significant impact on the mRNA expression levels of the glucocorticoid receptor gene (*Nr3c1*) in the medial prefrontal cortex, with adult rats exhibiting higher Nr3c1 mRNA levels relative to adolescent rats (diet: *F*_(1, 20)_ = .0002, *p* = .98; age: *F*_(1, 20)_ = 5.14, *p* = .03; interaction: *F*_(1, 20)_ = .01, *p* = .9) (**Figure 6B**). Although the adult rats showed a trend for higher mRNA levels of the glucocorticoid receptor chaperones, *Fkbp51* (Figure Not Shown) and *Fkbp52* (**Figure 6C**), analyses showed no significant effects of the diet type, age, and interactions between factors (for FKBP5-1, diet: *F*_(1, 20)_ = .34, *p* = .56; age: *F*_(1, 20)_ = 1.97, *p* = .17; interaction: *F*_(1, 20)_ = .05, *p* = .82; for FKBP5-2, diet: *F*_(1, 19)_ = .65, *p* = .42; age: *F*_(1, 19)_ = 3.60, *p* = .07; interaction: *F*_(1, 19)_ = 2.32, *p* = .14).

**Figure 6.**
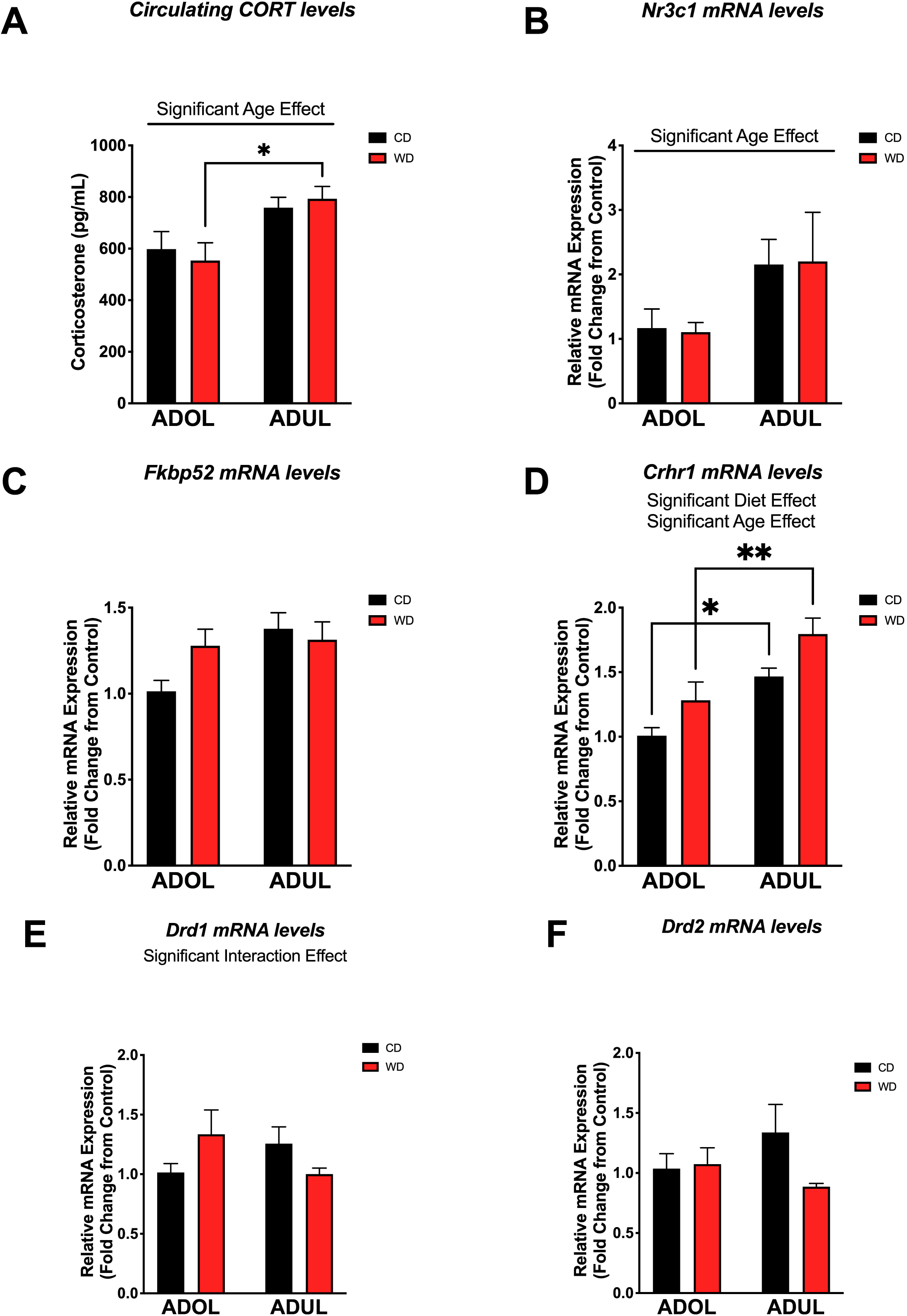
Age and short-term exposure to obesogenic diet increase stress biomarkers in the medial prefrontal cortex. **(A)** Age increased plasma corticosterone levels in WD rats (\**p* = .01; *n* = 6-8 rats/group). **(B)** Similarly, age had a significant impact on the messenger RNA levels of glucocorticoid receptor gene (*Nr3c1*) (\**p* = .03; *n* = 8 rats/group). **(C)** Age and diet had no effects on mRNA levels of FK506-binding protein 2 (*Fkpb52*) (*p* > .05; *n* = 8 rats/group). **(D)** Both age (*p* = .0002) and diet (*p* = .0093) had robust effects in the expression of the corticotropin-releasing hormone receptor 1 (*Crhr1*) mRNA. **(E)** There was a significant interaction between age and the obesogenic diet on the expression of the dopamine receptor D1 (*Drd1*) (*p* < .05; *n* = 7-8 rats/group). **(F)** There were no significant differences in the mRNA levels of the dopamine receptor D2 (*Drd2*) (*p* > .05; *n* = 7-8 rats/group). *, *p* < .05; **, *p* < .01. Data represents mean ± S.E.M.

The corticotropin-releasing hormone receptor 1 (*Crhr1*) is abundantly expressed in the mPFC and serves crucial roles in the regulation of stress and fear responses. We found that the rats that were exposed to the short-term obesogenic diet had higher Crhr1 mRNA levels in the mPFC relative to controls (diet: *F*_(1, 20)_ = 8.28, *p* = .009) (**Figure 6D**). Adult rats showed increased Crhr1 mRNA levels in the mPFC relative to adolescent rats (age: *F*_(1, 20)_ = 21.58, *p* = .0002; interaction: *F*_(1, 20)_ = .07, *p* = .79). Sidak’s post hoc analysis revealed that adult rats in both CD (*p* = .01) and WD (*p* = .004) groups had significantly higher Crhr1 mRNA levels in the PFC when compared to adolescent rats. Previous studies indicate the active involvement of dopaminergic receptors in diet-induced obesity and fear-related behaviors. Of interest, evidence indicates that dopamine receptors are under regulatory actions of CRH, and dictate fear conditioning and fear extinction responses. Thus, we aimed to determine the effects the obesogenic diet, age, and interactions on the mRNA levels of dopamine receptors 1 and 2. While diet [*F*_(1, 19)_ = .05, *p* = .82] and age [*F*_(1, 19)_ = .11, *p* = .73] did not have significant effects on the mRNA expression levels of the dopamine receptor 1 (*Drd1*), we found a significant interaction between these factors [*F*_(1, 19)_ = 4.39, *p* = .04] (**Figure 6E**). We found no significant effects of the diet [*F*_(1, 18)_ = 1.71, *p* = .20], age [*F*_(1, 18)_ = .13, *p* = .72], or interactions [*F*_(1, 18)_ = 2.38, *p* = .14] on the expression of Drd2 mRNA levels in the mPFC (**Figure 6F**).

## 5. DISCUSSION

Childhood trauma survivors who suffer from post-traumatic stress disorder (PTSD) are at high risk for developing obesity and metabolic disorders (Assari et al., 2016; Brewerton and ONeil, 2016; Llabre and Hadi, 2009; Ramirez and Milan, 2016; Roenholt et al., 2012). We reported evidence in support of these observations while demonstrating a novel directionality in the relationship between early-life posttraumatic stress reactivity and diet-induced obesity (DIO) (Kalyan-Masih et al., 2016; Vega-Torres et al., 2018). Our findings indicate that the consumption of an obesogenic Western-like high-saturated fat/high-sugar diet (WD) during adolescence heightens stress reactivity while altering key substrates implicated in PTSD (Kalyan-Masih et al., 2016; Vega-Torres et al., 2018).

Although the long-term impact of DIO on emotion regulation is becoming clear, mechanistic studies are confounded by the multiple complications associated with obesity. In this study, we expanded our previous observations and investigated the impact of a short exposure to a WD, including groups exposed to the WD during adolescence and adulthood. Importantly, we tested the acute WD effects, relative to an ingredient matched low-fat semi-purified control diet, on cued fear conditioning and fear extinction using the fear-potentiated startle (FPS) paradigm. The primary findings of this study are: **(a)** short exposure to a WD impairs cued fear extinction memory retention in adolescent rats, **(b)** short-term WD consumption attenuates basomedial amygdala activation to a fear conditioning paradigm, **(c)** short-term WD consumption increases CRHR1 mRNA expression levels in the medial prefrontal cortex, **(d)** adult rats exhibit markers of heightened HPA axis tone and emotional reactivity relative to adolescent rats. Altogether, this follow-up study provides supportive evidence that behavioral and molecular substrates implicated in fear are selectively vulnerable to the consumption of an obesogenic WD during adolescence. Our findings also serve to emphasize the caution that must be exercised when interpreting experimental outcomes when unmatched diets are used as controls in DIO studies.

### Short-term WD consumption attenuates fear extinction learning

An important finding of this study is that short-term WD consumption was sufficient to attenuate extinction learning of cued fear. Fear extinction has been characterized as an active form of inhibitory learning that allows for the adaptive regulation of conditioned fear responses (Myers and Davis, 2002). The inability to consolidate extinction memory and inhibit conditioned fear under safe conditions underlies some of the hallmark symptoms of anxiety and stress-related disorders (McGuire et al., 2016; Milad et al., 2009; Waters and Pine, 2016). It is now recognized that cognition, attention, mood, and anxiety disorders have a nutritional component or are promoted by poor dietary habits. Our results are consistent with evidence showing that the consumption of obesogenic high-fat diets can attenuate cognitive functions in humans and rodents in as little as 3-9 days (Beilharz et al., 2014; 2016; Edwards et al., 2011; Holloway et al., 2011; Kanoski and Davidson, 2010; Khazen et al., 2019; Sobesky et al., 2016). The findings of this study indicate that the adverse effects of consuming obesogenic foods expand to additional cognitive domains regulating emotional memories and fear. This is in agreement with new studies showing that adolescent rats that consume high-fat/high-sugar diets exhibit delayed spontaneous extinction and impaired extinction retention of fear-related behaviors (Baker et al., 2016; Reichelt et al., 2015; Vega-Torres et al., 2018). Our findings extend beyond those reported to date by showing fear extinction deficits independent of effects associated with obesity and metabolic disturbances.

Evidence supports that impairments in attention and threat discrimination may heighten the risk for anxiety and stress-related psychopathology (Block and Liberzon, 2016; López-Aumatell et al., 2009a; 2009b). This study confirms our previous findings showing that rats that consume obesogenic diets exhibit alterations in ASR plasticity, which may result in an inaccurate assessment of the level of threat (Kalyan-Masih et al., 2016; Vega-Torres et al., 2018). Notably, we found that the ASR neural substrates impacted by the WD seem to be dependent on the age of onset of diet exposure. The adolescent rats that consumed the WD exhibited increased ASR sensitization to the footshocks. This behavioral proxy may represent maladaptive stress reactivity and anxiety (Conti and Printz, 2003; Shalev et al., 2000).

Interestingly, the adult rats that consumed the WD showed greater long-term habituation of the ASR relative to CD rats. Since sensitization and habituation of the ASR require independent neural substrates (Koch, 1999; Leaton, 1976; Pilz and Leaton, 1999), our data indicate that short exposure to a WD influences different targets in adolescent and adult rats. Taken together, the ASR behavioral outcomes reported here support the notion that perturbations in attention, threat discrimination, and startle habituation may heighten vulnerability for anxiety and stress-related disorders in individuals that consume obesogenic diets.

### Short exposure to an obesogenic diet attenuates basomedial amygdala activation following cued fear conditioning

Several studies indicate that the medial prefrontal cortex (mPFC) and the basolateral complex of the amygdala (BLA) are critical to the acquisition and expression of conditioned fear (Davis, 2006; Quirk et al., 2006; Sotres-Bayon et al., 2004). The highly conserved neurocircuitry connecting the mPFC and the amygdala plays a critical role in anxiety, and in the extinction of fear memories (Adhikari et al., 2015; Janak and Tye, 2015; Milad and Quirk, 2002; Phelps et al., 2004) and is abnormal in PTSD patients (Gilboa et al., 2004; Koenigs and Grafman, 2009). The PFC and amygdala undergo striking structural changes during adolescence (Jalbrzikowski et al., 2017), providing a biological basis that may underlie their unique vulnerability to the disruptive effects of obesity and the consumption of diets rich in saturated fats and sugars. Paralleling clinical data in humans (Geha et al., 2017; Riederer et al., 2016), we showed that DIO rats exhibit significant and partly irreversible microstructural alterations in mPFC regions and amygdalar nuclei associated with fear learning and fear extinction (Vega-Torres et al., 2018). The impact of obesogenic diets on PFC and BLA neuroplasticity is supported by studies showing reduced dendritic spine density in the PFC (Dingess et al., 2017) and dendritic length in the basal arbors of the BLA (Janthakhin et al., 2017) in rats that consume obesogenic diets rich in fats. Together, these structural alterations may lead to anxiety (Kim and Whalen, 2009), impairments in fear processing (Poulos et al., 2009), and aberrant feeding behaviors (Land et al., 2014).

### Age increases anxiety and emotional reactivity in Lewis rats

There is some evidence in support of the notion that limbic structures involved in higher processing of emotional cues become deficiently activated with age despite showing a higher basal level of activation (Meyza et al., 2011). The histological results presented in this study provide support to this notion by indicating different c-Fos signatures in adolescent rats, with greater activation in mPFC and amygdalar regions, relative to adult rats. This suggests that the typical age-related increases in anxiety in adult LEW rats may be related to a blunted neuroendocrine response, combined with insufficient top-down regulation of limbic regions fear and anxiety (Adhikari et al., 2015).

It is becoming increasingly clear that obesogenic diets dysregulate critical mediators of hypothalamic-pituitary-adrenal (HPA) axis in rats (Abildgaard et al., 2014; Auvinen et al., 2012; Boitard et al., 2015; Kalyan-Masih et al., 2016; Khazen et al., 2019; McNeilly et al., 2015; Sobesky et al., 2016; Vega-Torres et al., 2018). The present findings seem difficult to reconcile with these reports as they indicate that short exposure to a WD was not sufficient to increase the circulating levels of corticosterone and related neuroendocrine biomarkers in the prefrontal cortex. However, differences in the methodology (fear conditioning paradigm) and diet composition (fat content and source; purified ingredient-matched control diet) may explain this discrepancy. Nonetheless, our findings support an age-related enhancement of the HPA axis tone in LEW rats (Meyza et al., 2011).

### Age and diet modulate the expression of the corticotropin-releasing hormone receptor 1

The corticotropin-releasing hormone (CRH) system has received considerable attention as a promising therapeutic target for anxiety and stress-related disorders (Bale and Vale, 2004; Binder and Nemeroff, 2010). CRH is an essential component of the behavioral and endocrine responses to stress, signaling through the stimulation of the G-protein-coupled CRH receptor 1 (CRHR1). The CRHR1 is abundantly expressed in fear-modulating corticolimbic circuits, including the mPFC (Steckler and Holsboer, 1999). Studies in humans and rodents indicate that CRH-CRHR1 hyper-signaling represents a candidate mechanism for PTSD risk (Bremner et al., 1997; Jovanovic et al., 2020; Rajbhandari et al., 2015; Toth et al., 2016). CRHR1 hyper-signaling alters brain structural integrity (Chen et al., 2004; Kolber et al., 2010; Toth et al., 2014) and has long-lasting consequences in stress susceptibility (Uribe-Mariño et al., 2016), mainly when it is triggered early in life (Toth et al., 2016).

Although the role of CRH in obesity is very complex, studies demonstrate that the CRH system has anorectic and thermogenic roles (Kuperman et al., 2016). CRH-CRHR1 signaling has emerged as a potential neuromodulator of food intake energy expenditure (Arase et al., 1988). Interestingly, CRHR1 is co-expressed with various metabolic receptors in corticolimbic structures (Koorneef et al., 2018), supporting an interplay between metabolism and fear. Our findings indicate that CRHR1 upregulation represents an early molecular adaptation to the obesogenic diet. Furthermore, our results provide support for further study of this signaling system as a candidate mechanism for anxiety and PTSD risk in obesity. We have demonstrated that exposure to an obesogenic diet during the critical maturational period of adolescence may permanently modify corticolimbic circuits and the response to further stressful stimuli. CRHR1 may play an integral role in the mechanisms of these long-lasting structural changes (Yang et al., 2015). Together, this study provides compelling evidence of the close relationship between energy homeostasis and the function of corticolimbic pathways modulating fear.

### Limitations and Future Studies

This study had some limitations to be addressed in future research. The results should be interpreted with caution, as results of this rat model do not necessarily directly translate to the human condition. Whereas our results are consistent with deficits in fear extinction, it is unclear from our data whether the WD effects are related to impairments in fear extinction memory acquisition, consolidation, expression, reconsolidation, and/or retrieval. Continued translational work will inform the basis of these learning and memory deficits. We analyzed mPFC markers only based on our prior studies revealing long-lasting effects of a short-term exposure to an obesogenic diet in this region and its critical role in fear extinction learning, retention, and expression (Vega-Torres et al., 2018). Future studies investigating the molecular landscape of the amygdala and other mesocorticolimbic structures would better characterize the impact of obesogenic diets in brain centers implicated with emotional regulation. Several lines of evidence demonstrate that high-fat diets and fear conditioning alter the HPA axis and dopamine function markers in the brain. Therefore, it is likely that a combination of these factors contributed to the observed differences in mRNA levels. Future studies are required to clarify the relative contribution of diet and fear conditioning to gene expression. Lastly, replication in different rat strains, gender, and conditioning paradigms is warranted.

### Summary

This study shows that the consumption of obesogenic diets during adolescence heightens behavioral vulnerabilities associated with risk for anxiety and stress-related disorders. Given that fear extinction promotes resilience against PTSD and fear extinction principles are the foundation of psychological treatments for anxiety and stress-related disorders, understanding how obesity and obesogenic diets affect the acquisition and expression of fear extinction memories is of tremendous clinical relevance.

## AUTHORS CONTRIBUTIONS

JDF and JDVT planned the experiments, tested data statistically and wrote the manuscript. JDVT conducted the behavioral and molecular experiments, including ASR, FPS, and EPM. MA and RRO performed the histological experiments and quantified c-Fos staining. ALRR contributed to the RT-qPCR studies, discussion, and the manuscript. POA conducted and analyzed the ELISA data.

## ACKNOWLEDGEMENTS

This study was partly supported by the NIH (P20MD006988 and 2R25 GM060507) and the Loma Linda University School of Medicine Seed Grant Funds to JDF. We would like to thank the staff at the animal care facility.

